# Functional and structural resilience of the active site loop in the evolution of *Plasmodium* lactate dehydrogenase

**DOI:** 10.1101/386029

**Authors:** Jacob D. Wirth, Jeffrey I. Boucher, Joseph R. Jacobowitz, Scott Classen, Douglas L. Theobald

## Abstract

The malarial pathogen *Plasmodium falciparum (Pf)* is a member of the Apicomplexa, which independently evolved a highly specific lactate dehydrogenase (LDH) from an ancestral malate dehydrogenase (MDH) via a five-residue insertion in a key active site loop. *Pf*LDH is widely considered an attractive drug target due to its unique active site. Apicomplexan loop conservation suggests that a particular insertion sequence was required to evolve LDH specificity, and we previously showed (Boucher 2014) that a tryptophan in the insertion, W107f, is essential for activity and specificity. However, the roles of other residues in the loop are currently unknown. Here we show that *Pf*LDH activity is remarkably resilient to radical perturbations of both loop identity and length. Thus, alternative insertions could have evolved LDH specificity as long as they contained a tryptophan in the proper location. *Pf*LDH therefore has high potential to develop resistance to drugs that target its distinctive active site.

## Introduction

Apicomplexa are obligate, intracellular eukaryotic parasites of animals responsible for many human diseases, including malaria, cryptosporidiosis, babesiosis, cystoisosporiasis, cyclosporiasis, and toxoplasmosis. The organisms responsible for malaria are of the genus *Plasmodium*, with *Plasmodium falciparum* responsible for the most lethal form of malaria. *P. falciparum* proceeds through a complex life-cycle in two different hosts. *P. falciparum* sporozoites infect humans *via* a mosquito bite. These sporozoites invade liver cells, where they multiply asexually into merozoites. The merozoites rupture the liver cells and enter the bloodstream, where they invade erythrocytes and replicate. The replication of *P. falciparum* in erythrocytes is known as the blood stage of infection, which is largely responsible for the clinical symptoms of malaria [1].

During the blood stage, *P. falciparum* must respire anaerobically to regenerate NAD^+^ [1, 2], an essential electron acceptor in glycolysis. In the presence of oxygen, ATP production is maximized by metabolizing glucose to CO_2_ and H_2_O *via* glycolysis, the citric acid cycle, and the electron transport chain, which regenerates NAD^+^ for use in glycolysis. In the absence of oxygen, NAD^+^ must be regenerated through pyruvate fermentation to prevent a stall in glycolysis and to generate sufficient amounts of ATP for cellular function. Lactate dehydrogenase (LDH) couples the regeneration of NAD^+^ from NADH to the reduction of pyruvate to lactate. Since the blood stage of malarial infection occurs under anaerobic conditions, the *Plasmodium falciparum* LDH (*Pf*LDH) is essential for the pathogen’s survival as the only means to regenerate NAD^+^ [2].

Apicomplexan LDHs evolved from an ancestral α-proteobacterial malate dehydrogenase (MDH), independently of canonical LDHs such as those found in the metazoan hosts of apicomplexan parasites [3]. Despite convergent evolutionary origins, *Pf*LDH and canonical LDHs both share a similar catalytic mechanism. The enzymatic reduction of pyruvate to lactate proceeds via the following steps: (1) starting in the “loop-open” conformation, NADH binds the *apo* enzyme, (2) R171 orients pyruvate in the active site to form the ternary complex, (3) the substrate specificity loop closes over the active site, allowing R109 to bind the substrate and stabilize the transition state, (4) NADH reduces pyruvate to lactate by hydride transfer, (5) the substrate specificity loop opens to release lactate, and (6) NAD^+^ is released to regenerate the *apo* enzyme. D168 activates the catalytic H195, allowing for a proton transfer during the reduction of pyruvate to lactate (**Figure 1a**). Movement of the substrate specificity loop is the rate-limiting step in catalysis [4].

**Figure 1:**
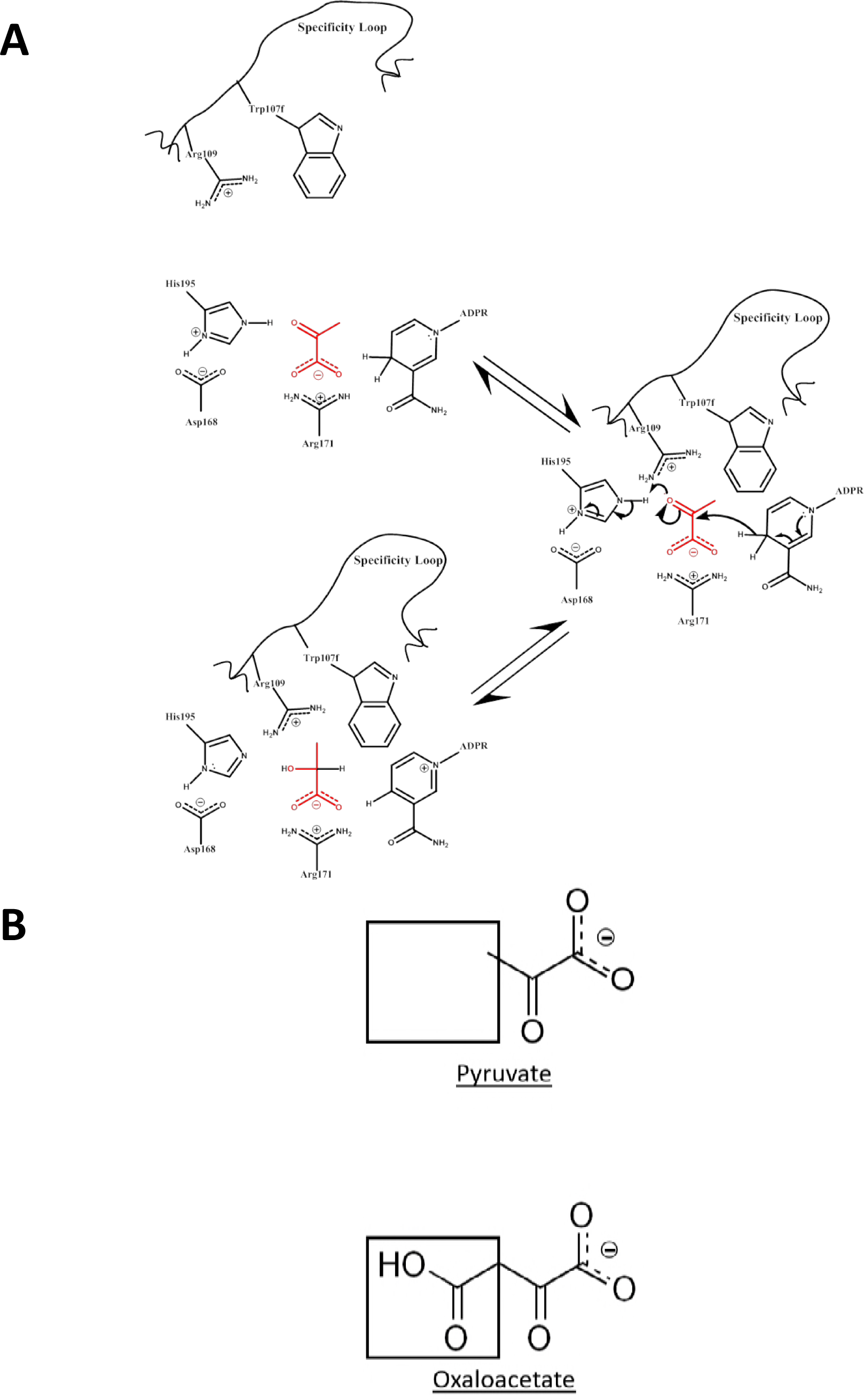
Mechanism of *Pf*LDH loop closure and catalysis. **(A)** NADH and pyruvate (**Red**) bind to the enzyme to form the ternary complex. Specificity loop closure brings Arg109 and Trp107f into proximity of the substrate and NADH donates a hydride to reduce pyruvate to lactate. **(B)** Black boxes outline the differing functional groups on pyruvate and oxaloacetate, a methyl or carboxylic acid group respectively.

The LDH substrate specificity loop also plays an important role in substrate recognition. This loop contains a residue at position 102, commonly referred to as the “specificity residue”, which distinguishes among the R-groups of different 2-ketoacid substrates. In all known MDHs, the specificity residue is an arginine, which forms a salt bridge with the methylene carboxylate group of malate and oxaloacetate (**Figure 1b**). In canonical LDHs the specificity residue is a glutamine, which contacts and recognizes the lactate and pyruvate methyl group (**Figure 1b**).

However, the convergent apicomplexan LDHs did not evolve by a mutation at position 102. Rather, apicomplexan LDHs evolved from an ancestral MDH via a unique five amino acid insertion in the substrate specificity loop that switches substrate specificity from malate/oxaloacetate to lactate/pyruvate [3]. The insertion lengthens the loop, but otherwise the structures of *Pf*LDH and canonical LDHs and MDHs are highly similar (**Figure 2a**).

**Figure 2:**
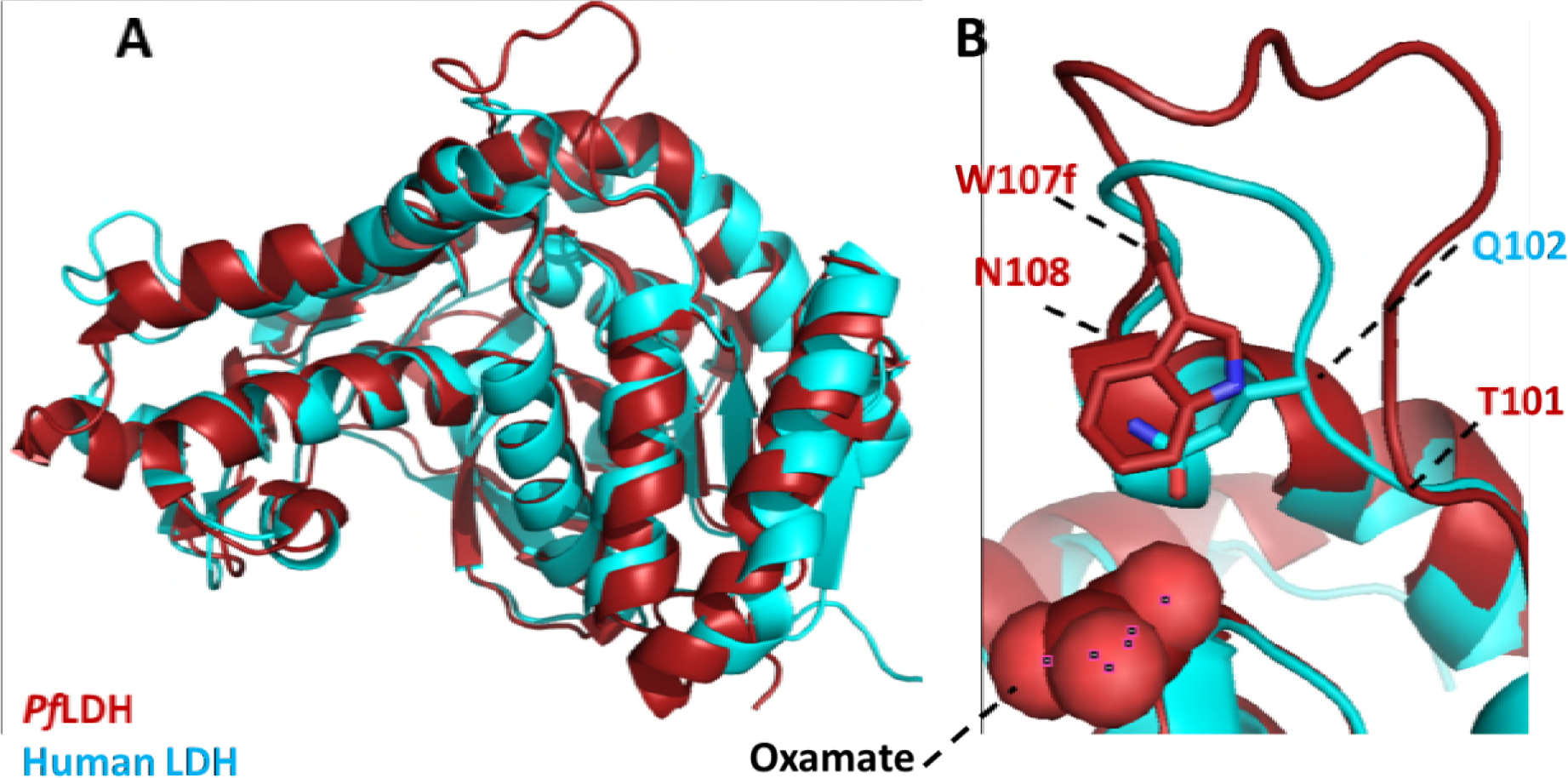
Superposition of Human (Teal; PDB: 1I10) and *P. falciparum* (Red; PDB: 1T2D) LDHs. **(A)** An ordinary least squares superposition of Human and *Pf*LDH (PyMol) shows high structural similarity. **(B)** A close-up of the loop shows the structural effect of the five amino acid insertion. *Pf*LDH’s W107f occupies the same three-dimensional space as the Human LDH’s Q102.

Crystal structures of canonical LDHs with the substrate specificity loop in the closed conformation show the Q102 specificity residue contacting substrate analogs (**Figure 2b**). In contrast, the *Pf*LDH crystal structure shows W107f contacting a substrate analog in the closed conformation, suggesting that W107f is the apicomplexan LDH “specificity residue” rather than the lysine at position 102 (**Figure 2b**). Using point mutations and alanine scanning of the substrate specificity loop, we previously showed that W107f is essential for *Pf*LDH enzyme activity and substrate recognition, while mutations of K102 have negligible effect [3].

*Pf*LDH is widely considered an attractive therapeutic target due to its essential role during pathogen replication in the blood stage and due to its distinct active site architecture, which provides an opportunity to selectively inhibit the parasite enzyme versus the host LDH. Development of small molecule inhibitors against *Pf*LDH is currently an active area of research [2, 5-13], and selective antibody inhibitors have been developed that target the unique *Pf*LDH substrate specificity loop [14]. Sequence comparison of apicomplexan LDHs suggests the specificity loop region is highly conserved (**Figure 3**). Hence, selective inhibitors of *Pf*LDH are also likely to be effective against various *Plasmodium* strains and other apicomplexan LDHs (for example, as with *Babesia* [15]).

**Figure 3:**
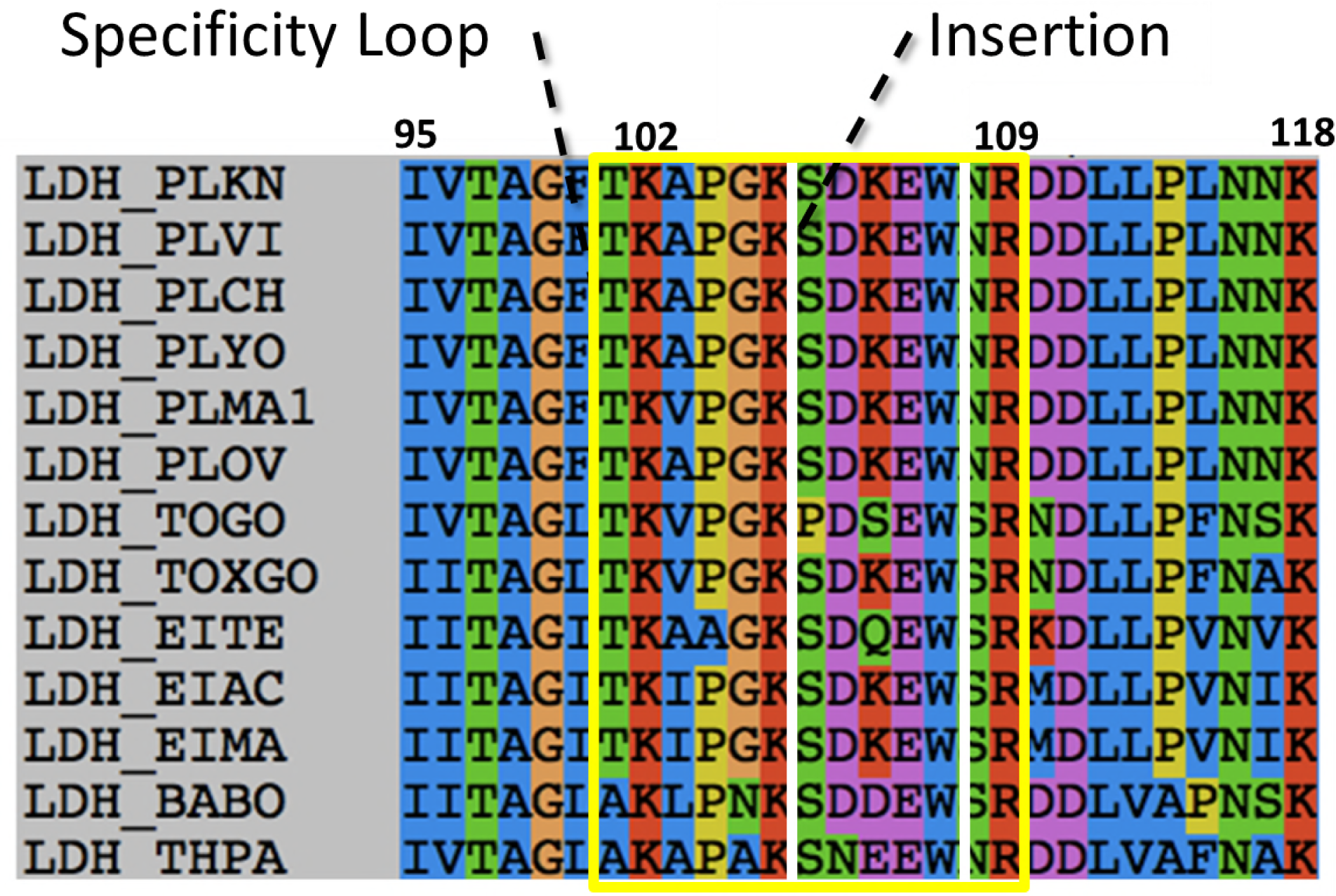
Multiple sequence alignment of the apicomplexan specificity loop. The loop is defined by positions 101 through 108 (as described in the text) and is boxed in yellow. The insertion found only in apicomplexan LDHs is boxed in white.

The apparent sequence conservation of the substrate specificity loop poses an evolutionary problem: Was the conserved insertion sequence necessary for evolution of LDH specificity from the ancestral MDH? Furthermore, crystal structures show that much of the apicomplexan LDH loop bulges out into solution, lacking any clear structural role without contacting the rest of the protein. Was the entire five-residue insertion necessary, or could a shorter insertion have sufficed? To answer these questions, we probed the functional constraints on residue identity and loop length within the *Pf*LDH substrate specificity loop by perturbation analysis. Loop conservation was quantified by estimating the evolutionary substitution rates across the loop, which identified several slow-evolving positions likely to be functionally constrained. With this information we generated an ensemble of *Pf*LDH loop mutants and characterized their steady-state kinetics. We also made a series of mutants with varying loop lengths to characterize the functional dependence on loop length. Surprisingly, neither sequence identity nor loop length are under strong functional constraint, as long as W107f is unperturbed.

## Methods

### Site-Directed Mutagenesis

The *Plasmodium falciparum* lactate dehydrogenase (LDH) gene (gi#: 124513226) was synthesized and subcloned into pET-11b by GenScript (Piscataway, NJ) with a C-terminal 6xHis-Tag. All mutations and truncations were generated using a QuikChange Lightning kit (Agilent, Santa Clara, CA) with primers synthesized by Integrated DNA Technologies (Coralville, IA). All sequences were confirmed by Sanger Sequencing at Genewiz (Cambridge, Massachusetts). Conventional LDH numbering is based on the canonical dogfish LDH sequence, which has no insertion [16]. The residues in the apicomplexan insertion are thus numbered using the residue that precedes the insertion, K107, with letters appended to each residue sequentially: G106, K107a, S107b, D107c, K107d, E107e, W107f, and N108.

### Protein Expression

Plasmids were transformed into BL21(DE3)pLysS *Escherichia coli* cells (Invitrogen, Grand Island, NY). Cells were grown at 37 °C and 225 RPM agitation in 2xYT media supplemented with 30mM potassium phosphate, pH 7.8 and 0.1% (w/v) glucose with cell growth monitored at OD_600_. When cultures reached OD_600_ 0.5-0.8, cells were induced with 0.5mM IPTG and incubated for four hours at 37 °C and 225 RPM agitation. Cells were pelleted by centrifugation and stored at −80°C.

### Purification

Cell pellets were resuspended on ice in 25mL of 50mM sodium phosphate, pH 8.0, 300mM sodium chloride, 10mM imidazole buffer and 375 units of Pierce universal nuclease (Thermo Scientific, Rockford, IL) per 1.5L of culture pellet. Resuspended cells were sonicated on ice for 2 minutes at 35% amplitude (30 seconds ON, 20 seconds OFF). Cell lysate was centrifuged for 20 minutes at 25,000xg to separate soluble and insoluble fractions. Supernatant was 0.22uM syringe filtered. The filtered lysate was purified by affinity chromatography with a 5mL HisTrap FF nickel column (GE Healthcare, Piscataway, NJ). Protein was eluted and fractionated with a 10mM to 500mM imidazole gradient. Fractions were inspected for protein content by SDS-PAGE, then pooled and concentrated in Amicon Ultracel-10K centrifugal filters (Millipore, Billerica, MA). Concentrated protein was buffer exchanged into 50mM tris, pH 7.4, 100mM sodium chloride, 0.1mM EDTA and 0.01% (w/v) azide *via* a PD-10 desalting column (GE Healthcare, Piscataway, NJ). The final enzyme concentrations were calculated by absorbance spectroscopy using Beer’s law with 280nm extinction coefficients and molecular weights given by ExPASy’s ProtParam program.

### Kinetic Assays

All reagents were purchased from Sigma-Aldrich (St. Louis, MO). Initial rates for the conversion of pyruvate to lactate were monitored on a CARY 100 Bio spectrophotometer (Agilent, Santa Clara, CA) by following the decrease in absorbance at 340nm associated with the conversion of NADH to NAD^+^. Kinetics assays were performed with variable pyruvate concentrations and constant NADH (200uM) in a 50mM tris, pH 7.5, 50mM potassium chloride buffer at 25 °C. The enzyme concentration ranged between 1uM and 1nM. Initial rates were plotted against substrate concentration and fit using KaleidaGraph to the Michaelis-Menten equation, v/[E]_t_ = k_cat_[S]/(K_M_ + [S]) or a substrate inhibition equation (v/[E]t = k_cat_ [S]/(K_M_ + [S] + [S]^2^/K_i_)). All assays were performed in triplicate, and errors reported as standard error of the mean (SEM, e.g. Supplemental Tables 1-5).

### Protein Crystallization, Data Collection, and Structure Determination

All reagents were used from Sigma-Aldrich (St. Louis, MO). Conditions were optimized based on promising conditions identified from screens with Crystal ScreenTM and Crystal Screen 2TM (Hampton Research, Aliso Viejo, CA). 4uL drops (2uL of 600uM or 20mg/mL protein stock and 2uL of well solution) were spotted on cover slips (Hampton Research, Aliso Viejo, CA) and equilibrated against 1mL of well solution by hanging-drop vapor diffusion at room temperature. Crystals were cryo-protected by soaking in 15% (w/v) dextrose solution for 3 minutes, transferring to a 30% (w/v) dextrose solution, and then immediately flash freezing the crystal in liquid N_2_.

Data sets were collected at the SIBYLS Beamline (12.3.1 Lawrence Berkeley National Laboratory, Berkeley, CA) and indexed, integrated, and scaled with XDS/XSCALE. PHENIX’s AutoMR program was used to solve the structures by molecular replacement with a previously solved *Pf*LDH model (PDB: 1T2D). Models were improved by rounds of manual refinement in Coot and automated refinement with PHENIX’s phenix.refine program. Model quality was determined using MolProbity in PHENIX. All structural alignments and images were generated using PyMOL.

### Evolutionary Conservation Estimation

Apicomplexan LDH sequences were collected from the NCBI database using the BLASTP searches with four query sequences, UniProt IDs: MDHC_PIG, Q76NM3_PLAF7, C6KT25_PLAF, and MDH_WOLPM for full coverage of the superfamily. Non-LDH Sequences were removed along with sequences shorter than 290 or longer than 340. The final dataset contained 277 sequences. A multiple sequence alignment of the dataset was generated using Muscle and analyzed using PhyML [17] and ConSurf [18]. The relative conservation of each amino acid position within the alignment was estimated by quantifying the relative substitution rate using Empirical Bayes maximum likelihood methods.

## Results

### Contributions from Individual Specificity Loop Residues

We previously used alanine scanning mutagenesis to assess the functional contribution of individual residues within the specificity loop [3]. The boundaries of the scan were determined from a superposition of the human lactate dehydrogenase (LDH) and *Pf*LDH (**Figure 2b**). The main chains of the specificity loops in the two structures conformationally diverge between positions 101 and 108, a span of twelve residues. These twelve residues were individually mutated to alanine in *Pf*LDH and characterized kinetically to evaluate the effects of each substitution on catalysis.

The alanine scan surprisingly revealed that all residues except W107fA tolerate mutation with little-to-no effect on their catalytic efficiency (k_cat_/K_M_, **Figure 4a**, as reported previously [3]). W107fA has a four order of magnitude drop in k_cat_, indicating that W107f is essential for wild-type levels of strong catalysis. Notably, k_cat_ remains relatively unaffected in all the other mutants (within a factor of two of the wild-type value). W107fA also has an increased K_M_, fifty times greater than wild-type. The remaining alanine mutants all have K_M_ values within an order of magnitude relative to wild-type *Pf*LDH (**Figure 4b**).

**Figure 4:**
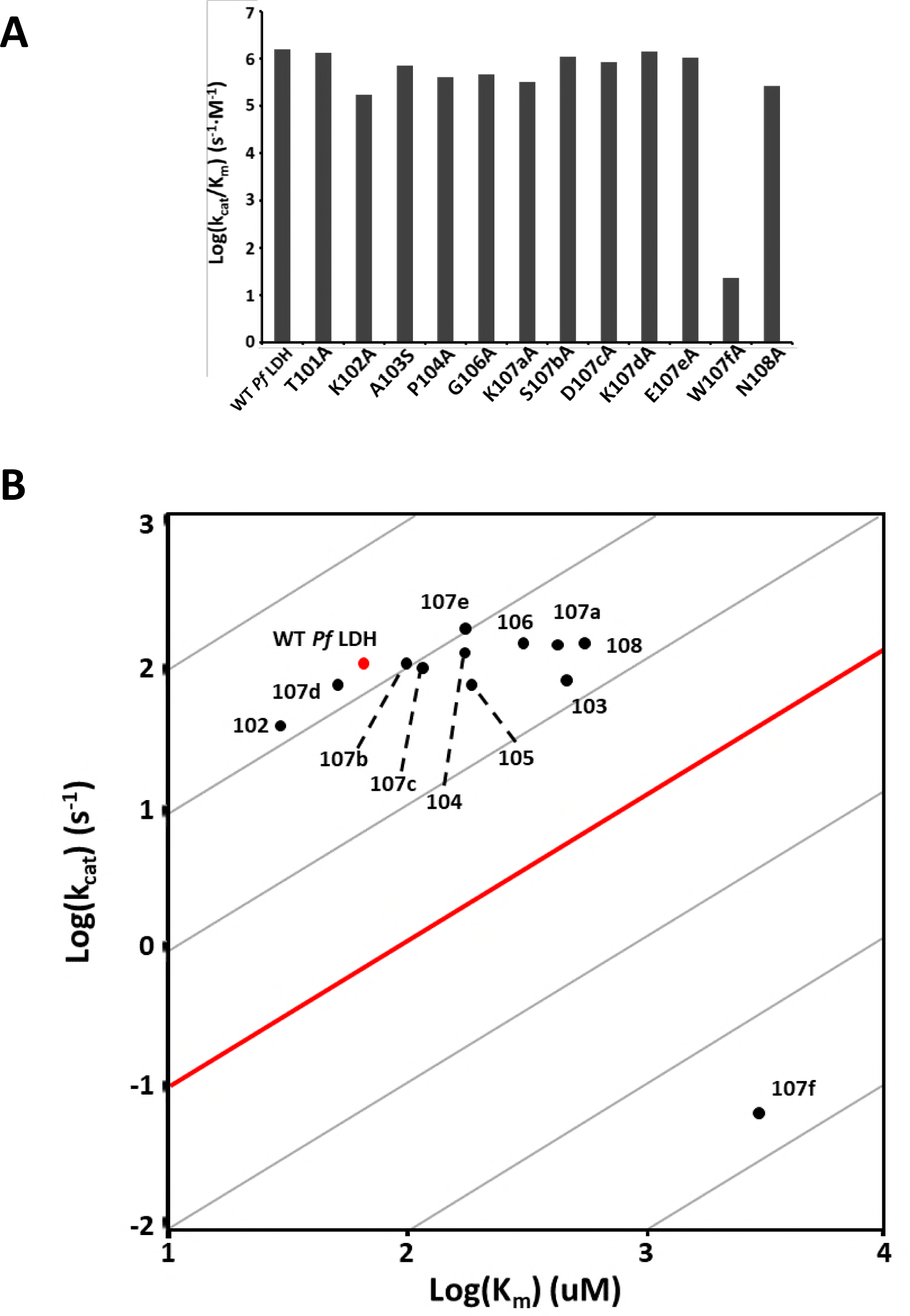
Alanine Scan k_cat_/K_M_ values, adapted and expanded from Boucher *et al* [3]. **(A)** Enzymatic activity for each mutant is displayed as log(k_cat_/K_M_). Substituting an alanine into any position in the loop other than 107f does not significantly affect pyruvate activity. SEM < 0.06 for all log(k_cat_/K_M_) values. **(B)** Kinetic parameters for each alanine mutant are plotted as log(k_cat_) vs log(K_M_). The red dot denotes wild-type *Pf*LDH and each black dot is labeled with the position that was mutated. The red line designates a k_cat_/K_M_ of 10^4^, roughly the minimum for physiologically relevant activity. Points on a grey line all have the same k_cat_/K_M_. W107fA has both a lower k_cat_ and higher K_M_ compared to wild-type. SEM < 0.02 for all log(k_cat_) values; SEM < 0.07 for all log(K_M_) values.

The W107fA crystal structure (PDBID: 4PLZ, Supp. Table 6) was solved by molecular replacement and was found to have the same space group and cell dimensions as the wild-type *Pf*LDH (PDBID: 1T2D) [3]. The W107fA structure is nearly identical to the wild-type protein (RMSD of 0.95 Å2, **Figure 5a**). The only significant difference between W107A and wild-type is in the orientation of the specificity loop. The wild-type loop is in the closed conformation, indicated by a kinked helix running along the back of the protein (**Figure 5b**). The W107fA mutant is in the open, non-catalytic conformation; indicated by the same helix in an unkinked linear conformation and disordered residues within the loop (**Figure 5b**).

**Figure 5:**
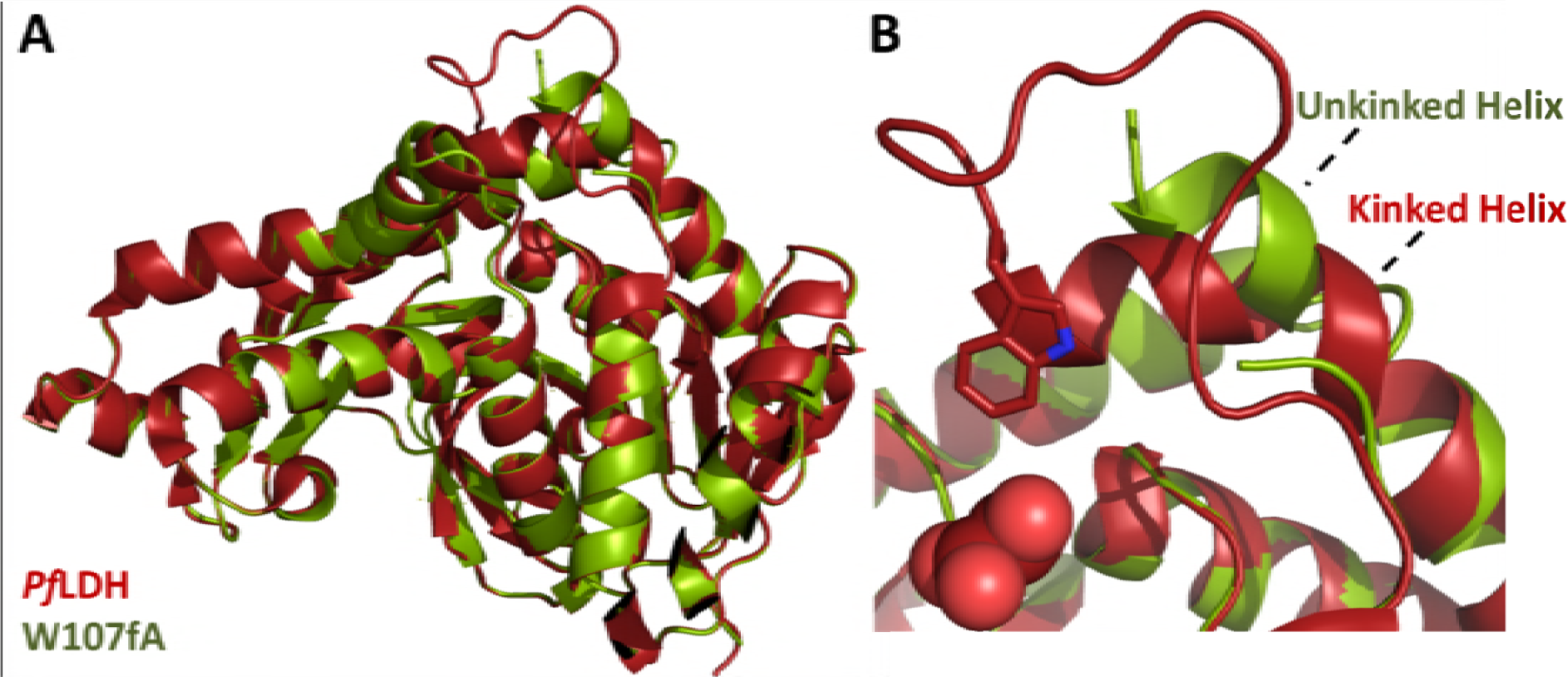
Loop conformation in the W107fA mutant. **(A)** Superposition of *Pf*LDH and the W107fA mutant (PyMol) indicates that the global LDH structure is essentially unperturbed by the substitution. **(B)** The mutant loop is in the open conformation and too dynamic or disordered to be modeled. This is the only significant difference in the two structures.

### Conservative W107f mutations

As mutating a tryptophan to an alanine is a dramatic mutation, we wished to investigate the tolerance of position 107f to more conservative changes. W107f was mutated to various hydrophobic and aromatic residues, in addition to a glutamine which is the canonical LDH specificity residue [19].

All W107f mutants had reduced enzymatic activity, with k_cat_/K_M_ values for pyruvate ranging from fifty times lower to five orders of magnitude below that of wild-type. Both k_cat_ and K_M_ for pyruvate are affected. The large aromatic replacements, W107fY and W107fF, have the least effect, with 50- and 100-fold reductions in pyruvate activity, respectively (**Figure 6a**).

**Figure 6:**
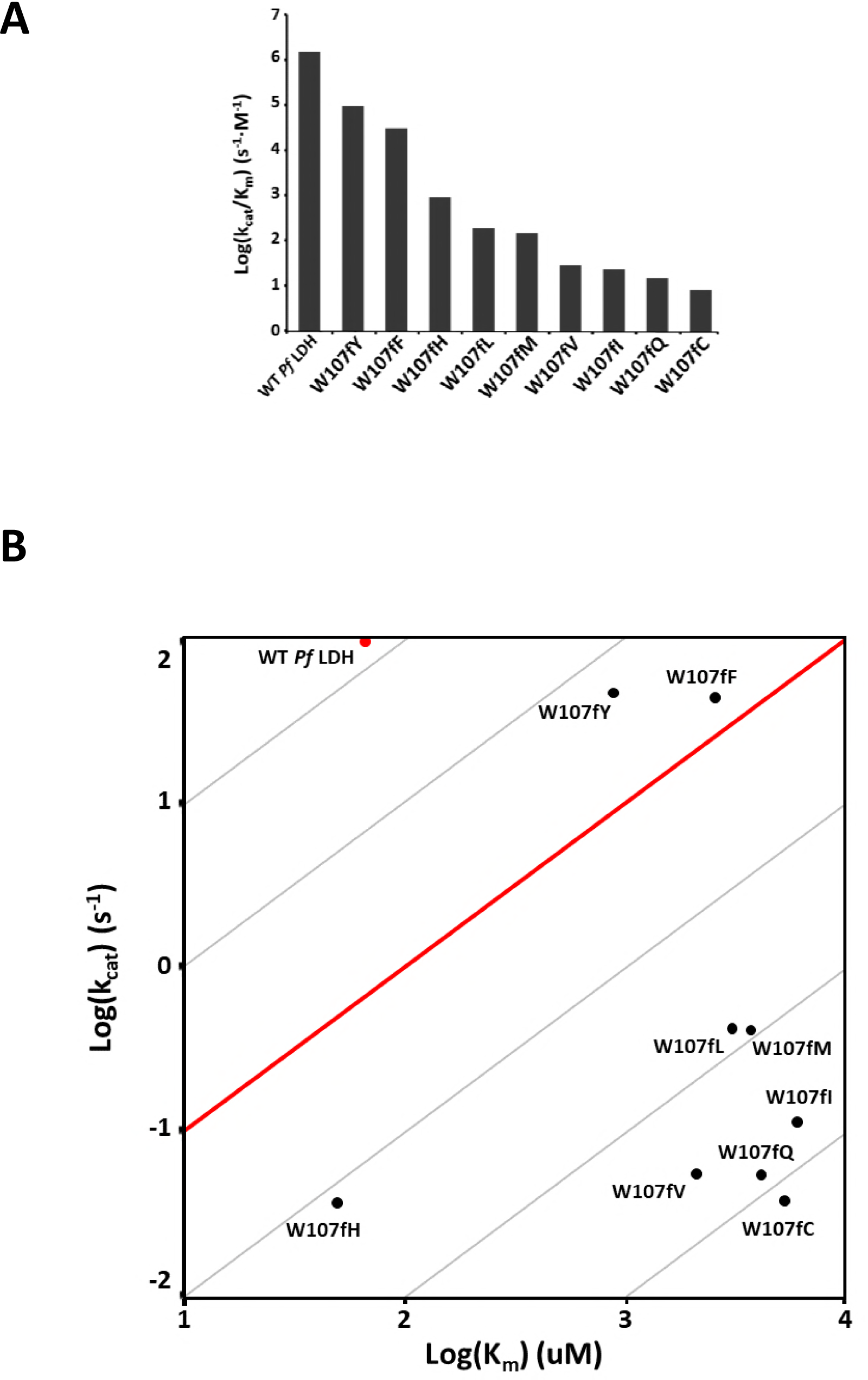
Kinetics of alternative W107f mutants. **(A)** Log(k_cat_/K_M_) values for more conservative W107f substitutions show that other residues at position 107f retain enzyme activity. All the constructs have levels lower than wild-type, yet the aromatic substitutions keep activity at physiological levels. SEM < 0.3 for all log(k_cat_/K_M_) values. **(B)** Analyzing the kinetic parameters separately shows that both tyrosine and phenylalanine have k_cat_ values nearly equal to wild-type *Pf*LDH, with the reduction in activity being primarily a K_M_ effect. Other mutants have both k_cat_ and K_M_ effects, except W107fH, which sees only a reduction in k_cat_. SEM < 0.1 for all log(k_cat_) values; SEM < 0.23 for all log(K_M_) values.

While these aromatic mutations have a substantial effect, the resulting activities are likely physiologically relevant as they are similar to those of other wild-type LDHs in the apicomplexan phylum, which have k_cat_/K_M_ values ranging from 10^4^ to 10^7^ [3, 20-26].

The other W107f mutants have more diminished activities. The smaller histidine, while still an aromatic residue, cripples the enzyme a further fifty times below W107fF. The non-planar hydrophobic amino acid mutants are all four orders of magnitude or more below the wild-type pyruvate k_cat_/K_M_, nearly on par with W107fA. Glutamine at position 107f is equally detrimental to pyruvate activity as the non-planar hydrophobics, despite being the specificity residue in canonical LDHs (**Figure 6a**). The substitutions have substantial effects on both the k_cat_ and K_M_ except for W107fY and W107fF, both of which have a k_cat_ within half of wild-type, and W107fH, which has a K_M_ nearly equal to wild-type (**Figure 6b**).

### Conservation of Loop Identity

Our alanine-scan mutants indicate most specificity loop residues are functionally unconstrained. This raises the question as to why the loop sequence is apparently well-conserved. High sequence conservation can arise for two main reasons, either (1) functional constraint to preserve the sequence or (2) insufficient time for the sequences to diverge. Phylogenetic rate analysis can control for evolutionary relatedness. Low rates indicate functional importance due to purifying selection for sequences identity, high rates suggest low functional constraint. Our rate analysis indicates K102, K107a, and E107e within the loop are relatively slow-evolving and likely functionally important (**Figure 7a**).

**Figure 7:**
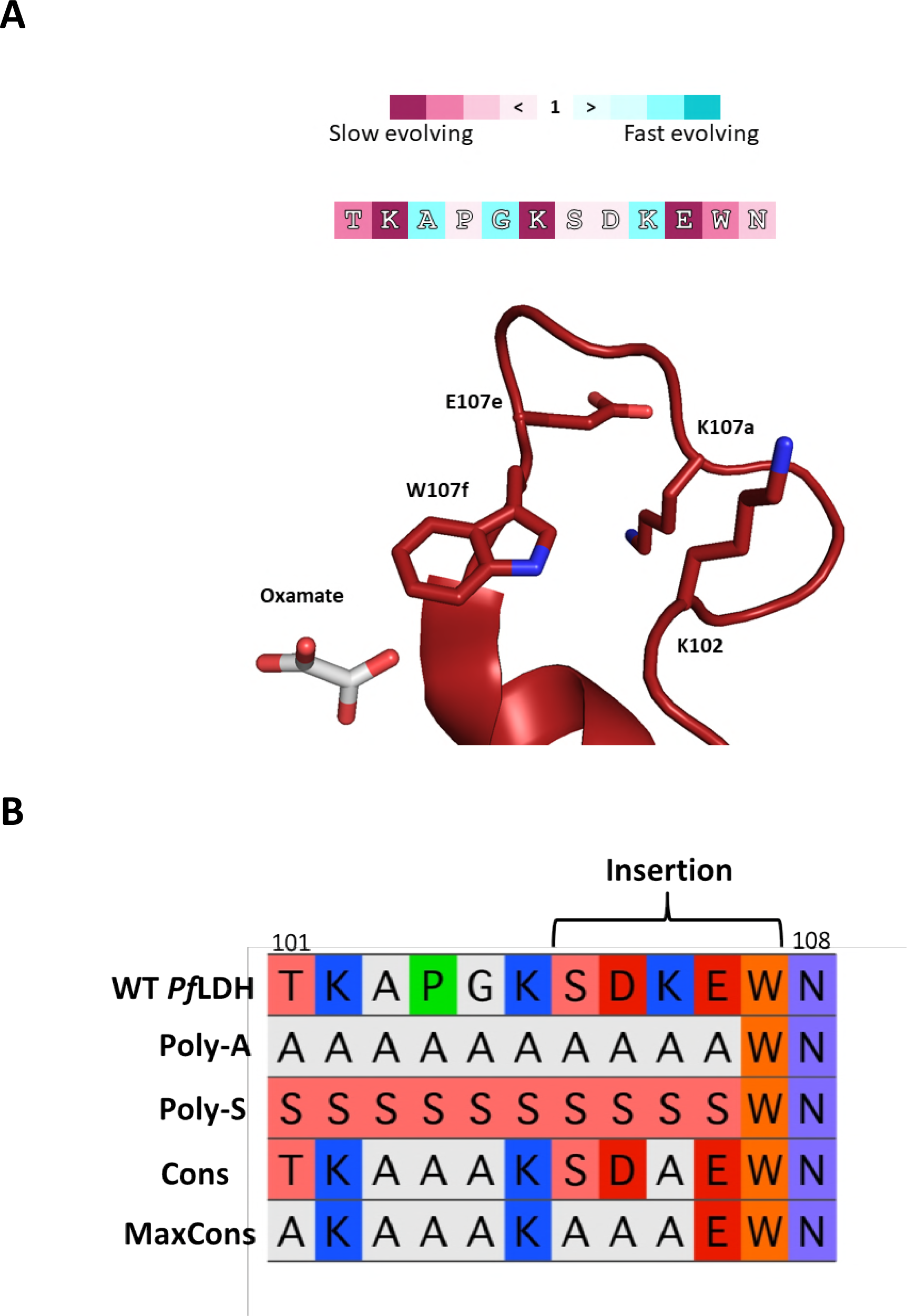
Evolutionary rates in the loop and conservation mutants. **(A)** Evolutionary rates are mapped onto the sequence of the *Pf*LDH specificity loop. Positions in magenta evolve slowly while those in cyan evolve more rapidly. Estimated rates are described in the methods. Positions with a rate less than 0.4 are shown as sticks in the *Pf*LDH crystal structure. **(B)** A multiple sequence alignment of the Poly-A, Poly-S, Cons, MaxCons mutants, and wild-type *Pf*LDH sequence.

We designed two *Pf*LDH mutant backgrounds to determine the role of the conserved residues. “Poly-A” and “Poly-S” mutants were constructed with all positions 101-107e mutated to alanine and serine, respectively (**Figure 7b**). The alanine scan indicates that any position other than W107f can be mutated individually with minimal effect on activity. Mutations in combination may have a more significant effect, and the Poly-A and Poly-S constructs test this by mutating each position simultaneously. The more polar Poly-S should provide a less drastic sequence change for a solvent-exposed surface loop and may indicate if changes in Poly-A function are due to the increased hydrophobicity. W107f was left unchanged in both mutant backgrounds because the alanine scan and the W107f substitution data show that the identity of position 107f is highly constrained. Remarkably, both the Poly-A and Poly-S mutants retain LDH activity within approximately two orders of magnitude of wild-type *Pf*LDH (**Figure 8**), a level of activity observed in other apicomplexan LDHs such as the *Toxoplasma gondii* LDHs [25].

**Figure 8:**
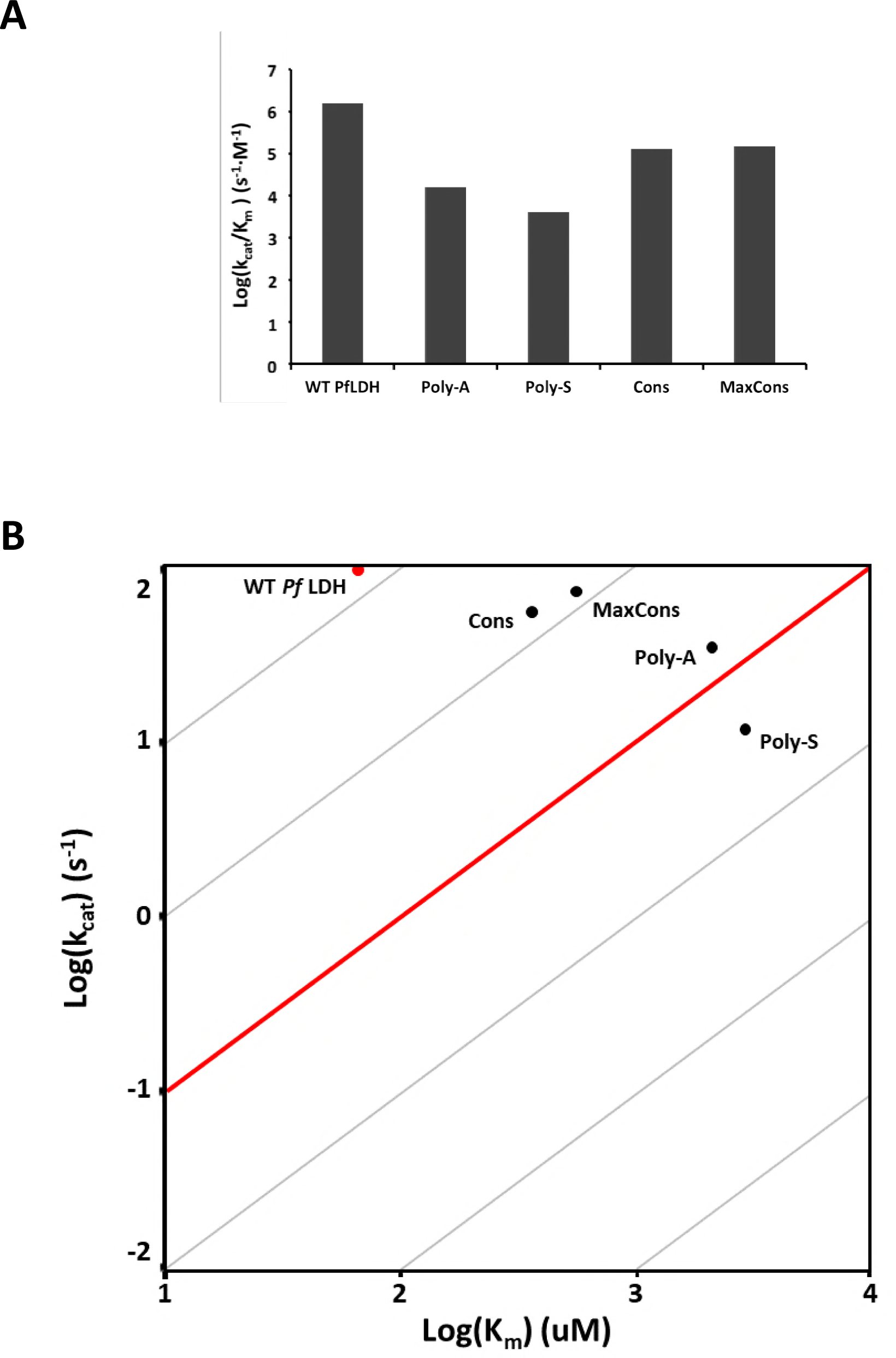
Activity of Poly-A, Poly-S, and conservation mutants. **(A)** Both “poly” mutants are reduced two orders of magnitude compared to wild-type. Re-addition of conserved residues in the Cons construct restores pyruvate activity 10-fold. The MaxCons construct shows that only K102, K107a, and E107e are necessary for the same level of activity. SEM < 0.03 for all log(k_cat_/K_M_) values. **(B)** The reintroduction of the highly conserved loop residues rescued activity both by increasing k_cat_ and reducing K_M_. MaxCons, the most active, shows a k_cat_ half that of wild-type *Pf*LDH and a K_M_ approximately 5-fold higher. SEM < 0.006 for all log(k_cat_) values; SEM < 0.032 for all log(K_M_) values.

We designed two additional mutants within the Poly-A background to determine the contributions from slow-evolving positions. If the loop residues are conserved due to functional constraint, there should be a partial rescue of activity when adding them back into the Poly-A mutant. The first construct “Cons” contains *Pf*LDH wild-type residues at any loop position evolving slowly (defined as relative rate <1): T101, K102, P104, K107a, S107b, D107c, and E107e (**Figure 7b**). Adding back the conserved residues partially restored activity by a factor of ten increase in k_cat_/K_M_ compared to Poly-A (**Figure 8)** but is still ten-fold less than wild-type *Pf*LDH activity. The second mutant “MaxCons” contains only the wild-type residues position that evolve as slowly as W107f (relative rate <0.4): K102, K107a, and E107e (**Figure 7b**). MaxCons behaved similarly to Cons despite having three fewer wild-type residues, suggesting that K102, K107a, E107e, and W107f are the most functionally important loop residues (**Figure 8**).

### Tolerance of the Specificity Loop to Truncation

The five-residue loop insertion is unique to the apicomplexan LDHs, which makes their specificity loop longer than what is seen in canonical LDHs [19]. The existence of highly active canonical LDHs with shorter specificity loops raises the question of whether the large *Pf*LDH loop is functionally necessary. Can *Pf*LDH retain activity with a shortened specificity loop? We therefore designed and tested a series of loop deletion mutants, ranging from one to five residues, to assess the dependence of *Pf*LDH activity on loop length (**Figure 9)**. Loop deletion mutants are named in the form of M(X) where X is the single letter amino acid code for the missing residues. Additionally, the “GSA-Linker” construct incorporates a generic glycine-serine-alanine motif that connects T101 to W107f, making the shortest loop possible that retains W107f (based on the distance between the backbone Cαs of T101 and W107f). GSA-Linker is effectively a 6-residue deletion.

**Figure 9:**
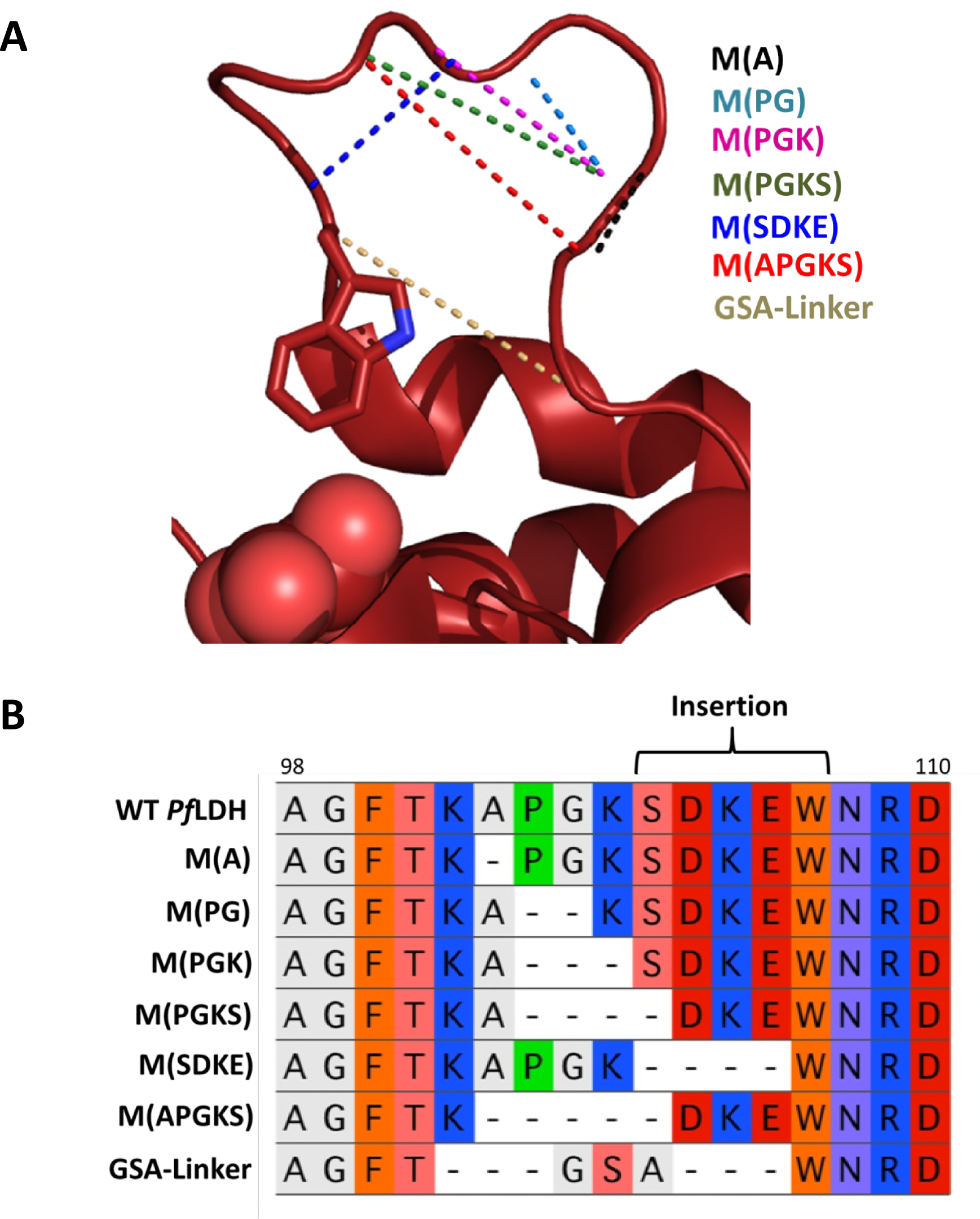
Loop truncation mutants. **(A)** A structural view of the specificity loop shows the connected residues of ligated positions in truncation mutants with dashed lines. The legend on the right color-coordinates each mutant to the dashed lines. **(B)** A protein sequence alignment of the truncation mutants. Dashes represent deleted amino acids.

M(A), M(PG), M(PGK), exhibit between a 10-fold and 100-fold reduction in k_cat_/K_M_ relative to wild-type (**Figure 10a**), within the range of other wild-type LDH activities in the apicomplexan phylum. The M(PGKS) and M(SDKE) truncations are more deleterious but remain marginally active. M(APGKS) and the GSA-Linker are crippled enzymes, with k_cat_/K_M_ values that are reduced approximately five orders of magnitude compared to wild-type.

**Figure 10:**
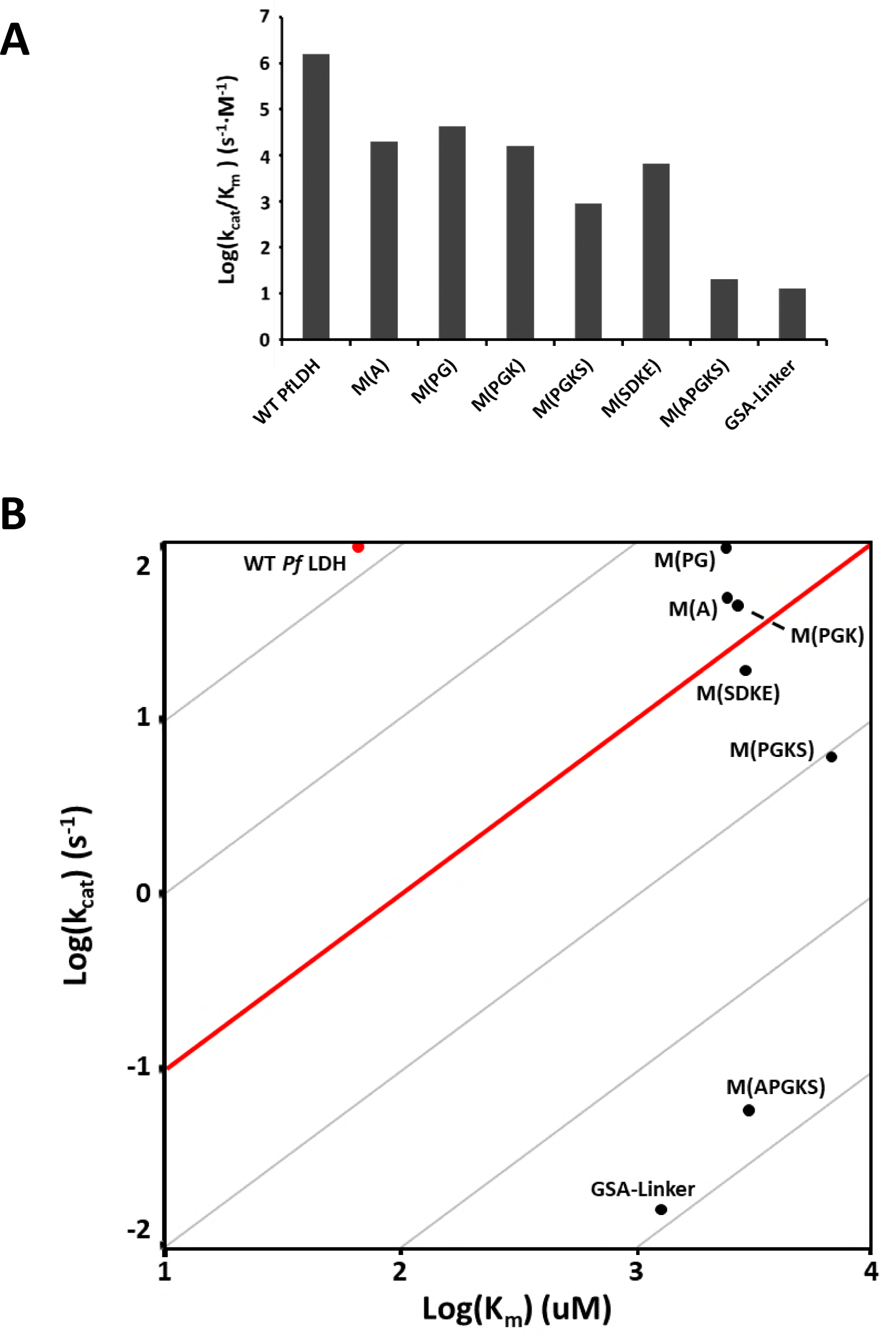
Kinetics of loop length mutants. **(A)** Truncation mutants all have reduced log(k_cat_/K_M_) when compared to wild-type. Deleting three loop residues or less reduces activity two orders of magnitude or less. Removing four residues has a more pronounced effect. The five-residue truncation and GSA-Linker are severely crippled. SEM < 0.06 for all log(k_cat_/K_M_) values except M(APGKS) and GSA-linker which have log(k_cat_/K_M_) SEM < 0.2. **(B)** The reduction in k_cat_/K_M_ for all the constructs is primarily a K_M_ effect, though both M(APGKS) and the GSA-linker have a significantly reduced k_cat_. SEM < 0.005 for all log(k_cat_) values except M(APGKS) and GSA-linker which have log(k_cat_) SEM < 0.2; SEM < 0.064 for all log(K_M_) values.

The loop deletions primarily affect K_M_, with only minor effects on k_cat_. One-to three-residue deletions reduce k_cat_ to no less than half of wild-type *Pf*LDH and both four-residue deletions reduce k_cat_ by an order of magnitude or less. Only the M(APGKS) and GSA-linker mutants have large reductions in k_cat_. However, all K_M_s of deletion mutants are thirty to forty times greater compared to wild-type, independent of the number of residues deleted (**Figure 10b**).

Deleting three residues appears to be the largest loop truncation that is still well tolerated, with a k_cat_/K_M_ > 10^4^. We next determined the kinetic effects of the position of the deleted residues in the loop by constructing and characterizing three-consecutive-residue loop truncations in all possible registers. All three-residue truncations had similar kinetics, with approximately a 100-fold reduction in k_cat_/K_M_, relative to wild-type, yet still within physiologically relevant range of activity (**Figure 11**).

**Figure 11:**
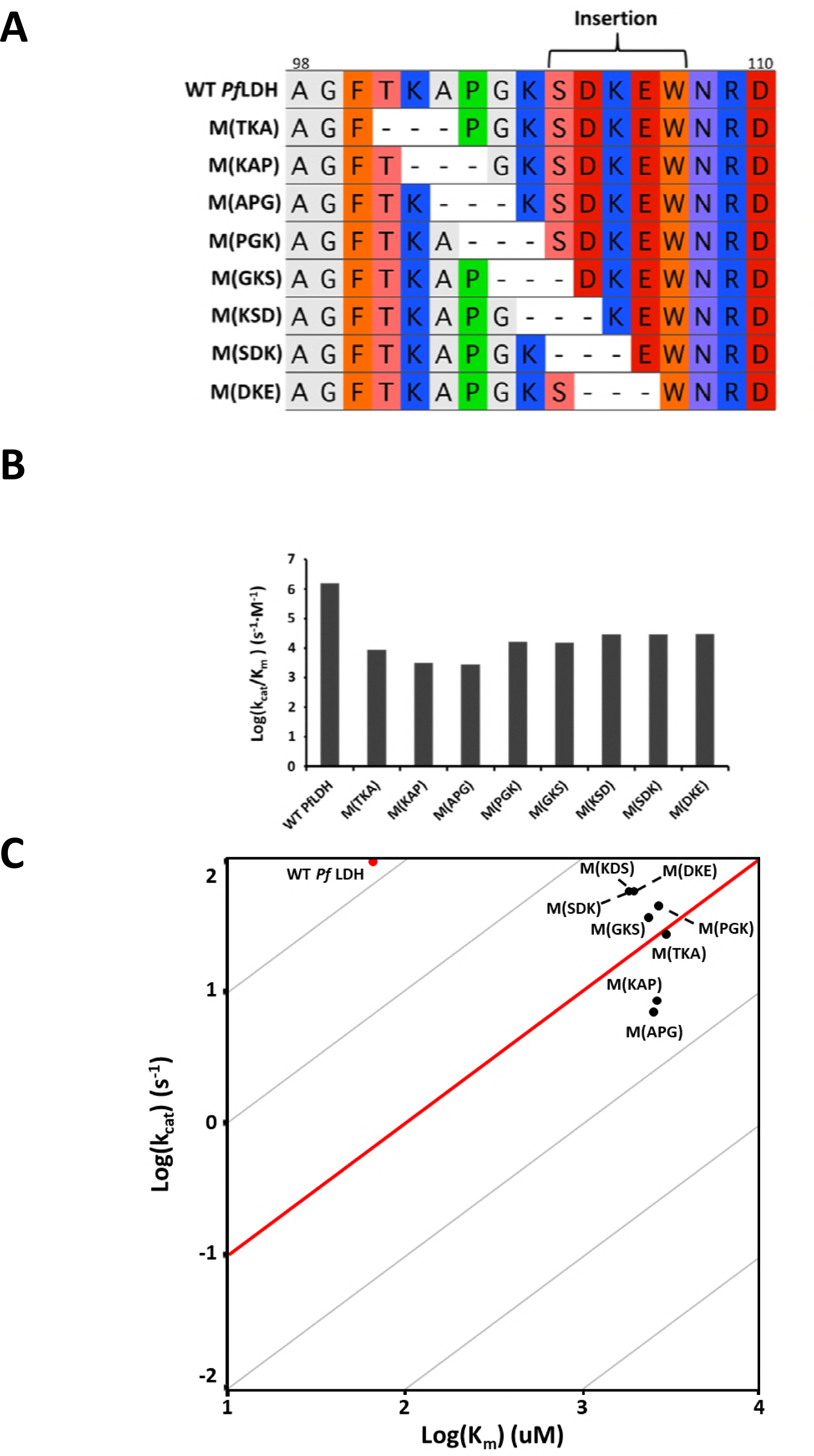
Positional effects of three-residue deletions. **(A)** A sequence alignment of the three-residue deletion mutants. Dashed indicates location of the gap. **(B)** All three-residue deletions are approximately one hundredth of the wild-type catalytic efficiency, regardless of the residues removed. SEM < 0.07 for all log(k_cat_/K_M_) values. **(C)** The majority of the activity loss is due to a decreased K_M_, which is approximately the same for all three-residue deletion mutants. SEM < 0.023 for all log(k_cat_) values; SEM < 0.08 for all log(K_M_) values.

## Discussion

### W107f is Under Extreme Functional Constraint

Alanine-scanning mutagenesis of the specificity loop revealed that only W107f contributes significantly to LDH activity (**Figure 4**). Additionally, most of the alternative residues at the 107f position fail to replicate wild-type levels of activity. Only the other large aromatic residues, tyrosine and phenylalanine retain physiologically relevant activity, though still below wild-type *Pf*LDH. (**Figure 6**). These results show that only by mutating a large planar aromatic residue at the 107f position could have conferred LDH activity to the apicomplexan ancestral MDH, but without needing to be tryptophan.

Unlike the wild-type, the crystal structure of W107fA is in the open conformation, suggesting W107f contributes favorable interactions necessary for loop closure as the W107fA mutant apparently prefers the open state (**Figure 5**). The rate-limiting step in pyruvate turn-over by *Pf*LDH is loop closure [4], and the reduction in k_cat_/K_M_ for W107fA results primarily from a decrease in k_cat_. Taken together, these facts suggest that k_cat_ of the W107fA mutant decreased because loop closure is slower. Given that K_M_ = (k_off_+k_cat_)/k_on_, and the difference in mutant and wild-type K_M_ is much smaller than the k_cat_ effect, there must be some compensatory change in the binding constants of the substrate: either k_off_ increased or k_on_ decreased. We propose that loss of favorable interactions from W107f result in a loop that prefers to remain open in a conformation unfavorable for substrate binding, and thus substrate is ejected before hydride transfer takes place.

### Conservation and Function Contribution of Specificity Loop Residues

Six of the ten loop residues evolve slower than average and thus may be functionally important (**Figure 7a**). The contribution of the three slowest evolving (K102, K107a, and E107e) is however modest, a 10-fold increase in k_cat_/K_M_ compared to a loop consisting entirely of alanines and W107f (**Figure 8**). Activity is not further rescued by incorporating additional slow-evolving wild-type residues.

Overall, *Pf*LDH is roughly 50% sequence identical to LDHs from closely related Apicomplexa, such as *Eimeria maxima* and *Toxoplasma gondii* [3, 25, 27, 28], yet the specificity loop region appears well-conserved. However, we find no obvious reason that the sequence and length of the specificity loop should be conserved, and in fact quantitive rate analysis indicates that most of the loop is not particularly slowly evolving. While the most conserved residues appear to contribute minimally to LDH function, it is possible the loop is important for another, unknown function. The loop sequence identity could be pleiotropically constrained if the enzyme is “moonlighting” [29, 30] and performs multiple functions for the organism (as seen in some vertebrate LDHs [31, 32]).

### The Specificity Loop can be Dramatically Truncated

The *Pf*LDH specificity loop can be truncated by as many as four residues and still maintain significant levels of activity (**Figure 10**). K_M_ is affected by loop truncations more than k_cat_, with all truncation mutants exhibiting a K_M_ an order of magnitude greater than wild-type, suggesting impaired substrate binding. Assuming k_on_ is diffusion limited and is unlikely to be affected, the loop truncations effects are probably mediated by an increase in substrate k_off_ due to a moderately unfavorable truncated loop conformation.

Loop deletion mutants may expose the active site to bulk solvent and increase the frequency of nonproductive binding events, which may explain why each of the three-residue deletion mutants all have approximately equal k_cat_/K_M_ regardless of the location in the specificity loop or the identity of the residues removed. The size of substrate loop is more important than loop identity during catalysis. However, there is a lower limit to how short the loop can be to remain functional.

### Evolutionary Requisites and Implications for Drug Resistance

Our results show that *Pf*LDH activity is extremely robust to mutations in the specificity loop that alter either its sequence identity or length. In fact, only W107f appears to be particularly important for enzyme function, and yet other aromatic residues can function nearly as well tryptophan. Ten of eleven possible positions are mutated in the Poly-A and Poly-S constructs and both are functional enzymes with physiologically relevant activity. Deletions of one, two, and three amino acids throughout the loop result in functional enzymes. These results suggest that the exact insertion sequence and length was unnecessary to evolve a novel LDH from the ancestral MDH. Many possible loop insertions could have evolved a functional LDH, regardless of sequence identity or length, with the only necessity being an aromatic amino acid properly positioned in the active site.

Drug resistance can develop if the protein target is not functionally constrained. There are many possible specificity loops that can produce physiologically relevant LDH activities in *P. falciparum*. *Pf*LDH thus has high potential to evolve mutations that could abolish any inhibitory effect of a drug that targets the active site loop because of the lack of significant functional constraint. The specificity loop region is conserved in the phylum and thus targeting the specificity loop in other Apicomplexa would likely be met with the same difficulties.

## Supplemental Materials

**Supplemental Table 1.** (Figure 4 Supp) Alanine Scan Kinetics Source Data

| Enzyme | $k_{\text{cat}} \pm \text{SEM} (\text{s}^{-1})$ | $K_{\text{M}} \pm \text{SEM} (\mu\text{M})$ | $k_{\text{cat}}/K_{\text{M}} (\text{s}^{-1} \cdot \text{M}^{-1})$ |
| --- | --- | --- | --- |
| WT PflLDH | $106 \pm 2$ | $67 \pm 5$ | $1.58 \times 10^6$ |
| T102A | $30 \pm 1$ | $30 \pm 4$ | $1.33 \times 10^6$ |
| K103A | $79 \pm 2$ | $463 \pm 57$ | $1.71 \times 10^5$ |
| A104S | $125 \pm 2$ | $176 \pm 13$ | $7.11 \times 10^5$ |
| P105A | $77 \pm 2$ | $190 \pm 25$ | $4.06 \times 10^5$ |
| G106A | $144 \pm 2$ | $312 \pm 22$ | $4.62 \times 10^5$ |
| K107aA | $141 \pm 2$ | $438 \pm 43$ | $3.21 \times 10^5$ |
| S107bA | $110 \pm 5$ | $100 \pm 16$ | $1.10 \times 10^6$ |
| D107cA | $98 \pm 4$ | $117 \pm 18$ | $8.38 \times 10^5$ |
| K107dA | $74 \pm 2$ | $52 \pm 6$ | $1.43 \times 10^6$ |
| E107eA | $185 \pm 7$ | $178 \pm 24$ | $1.04 \times 10^6$ |
| W107fA | $0.07 \pm 0.002$ | $3048 \pm 320$ | $2.30 \times 10^1$ |
| N108A | $147 \pm 4$ | $551 \pm 64$ | $2.67 \times 10^5$ |

**Supplemental Table 2.** (Figure 6 Supp) W107f Substitution Kinetics Source Data

| Enzyme | $k_{\text{cat}} \pm \text{SEM} (\text{s}^{-1})$ | $K_{\text{M}} \pm \text{SEM} (\mu\text{M})$ | $k_{\text{cat}}/K_{\text{M}} (\text{s}^{-1} \cdot \text{M}^{-1})$ |
| --- | --- | --- | --- |
| WT PflLDH | $106 \pm 2$ | $67 \pm 5$ | $1.58 \times 10^6$ |
| W107Y | $58 \pm 0.95$ | $870 \pm 44$ | $6.62 \times 10^4$ |
| W107F | $54 \pm 1.44$ | $2500 \pm 184$ | $2.18 \times 10^4$ |
| W107H | $0.04 \pm 0.01$ | $50 \pm 11$ | $7.40 \times 10^2$ |
| W107L | $0.48 \pm 0.02$ | $2975 \pm 356$ | $1.61 \times 10^2$ |
| W107M | $0.47 \pm 0.01$ | $3630 \pm 402$ | $1.29 \times 10^2$ |
| W107V | $0.06 \pm 0.01$ | $2066 \pm 1010$ | $2.66 \times 10^1$ |
| W107I | $0.13 \pm 0.01$ | $5837 \pm 881$ | $2.15 \times 10^1$ |
| W107Q | $0.06 \pm 0.01$ | $4020 \pm 369$ | $1.42 \times 10^1$ |
| W107C | $0.04 \pm 0.01$ | $5214 \pm 575$ | $7.94 \times 10^0$ |

**Supplemental Table 3.** (Figure 8 Supp) Loop Conservation Mutant Kinetics Source Data

| Enzyme | $k_{\text{cat}} \pm \text{SEM} (\text{s}^{-1})$ | $K_{\text{M}} \pm \text{SEM} (\mu\text{M})$ | $k_{\text{cat}}/K_{\text{M}} (\text{s}^{-1} \cdot \text{M}^{-1})$ |
| --- | --- | --- | --- |
| WT PflLDH | $106 \pm 2$ | $67 \pm 5$ | $1.58 \times 10^6$ |
| Poly-A | $34 \pm 0.8$ | $2090 \pm 162$ | $1.60 \times 10^4$ |
| Poly-S | $12 \pm 0.3$ | $2890 \pm 237$ | $4.10 \times 10^3$ |
| Cons | $71 \pm 0.8$ | $551 \pm 25$ | $1.30 \times 10^5$ |
| MaxCons | $54 \pm 0.8$ | $356 \pm 22$ | $1.50 \times 10^5$ |

**Supplemental Table 4.**
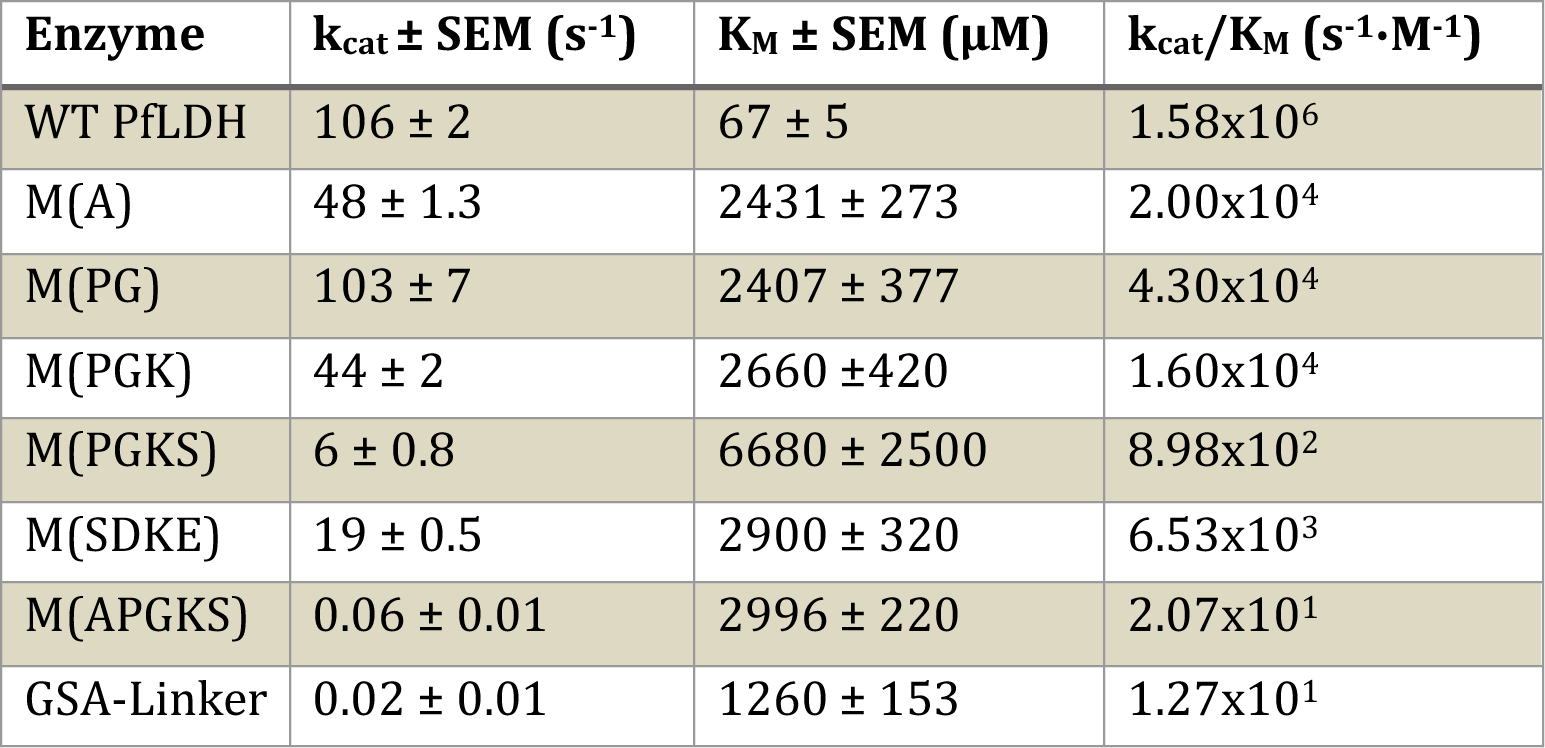
(Figure 10 Supp) Loop Truncation Mutant Kinetics Source Data

**Supplemental Table 5.** (Figure 11 Supp) Scanning Truncation Mutant Kinetics Source Data

| Enzyme | $k_{\text{cat}} \pm \text{SEM} (\text{s}^{-1})$ | $K_{\text{M}} \pm \text{SEM} (\mu\text{M})$ | $k_{\text{cat}}/K_{\text{M}} (\text{s}^{-1} \cdot \text{M}^{-1})$ |
| --- | --- | --- | --- |
| WT PflLDH | $106 \pm 2$ | $67 \pm 5$ | $1.58 \times 10^6$ |
| M(TKA) | $27 \pm 0.6$ | $3029 \pm 199$ | $8.90 \times 10^3$ |
| M(KAP) | $8.5 \pm 0.2$ | $2656 \pm 267$ | $3.20 \times 10^3$ |
| M(APG) | $7.0 \pm 0.1$ | $2537 \pm 199$ | $2.80 \times 10^3$ |
| M(PGK) | $44 \pm 2$ | $2660 \pm 420$ | $1.60 \times 10^4$ |
| M(GKS) | $36 \pm 0.8$ | $2365 \pm 192$ | $1.50 \times 10^4$ |
| M(KSD) | $56 \pm 2.9$ | $1952 \pm 374$ | $2.90 \times 10^4$ |
| M(SDK) | $56 \pm 2.9$ | $1952 \pm 374$ | $2.90 \times 10^4$ |
| M(DKE) | $56 \pm 1.9$ | $1862 \pm 240$ | $3.00 \times 10^4$ |

**Supplemental Table 6.** (Figure 5 Supp) Crystallographic Statistics for *Pf*LDH-W107fA

| **Data Collection** | *Pf*LDH-W107fA (4PLZ) |
| --- | --- |
| Space group | I 2 2 2 |
| Cell dimensions |  |
| *a*, *b*, *c* (Å) | 80.5, 86.0, 91.4 |
| $\alpha$, $\beta$, $\gamma$ (°) | 90, 90, 90 |
| Resolution (high res. bin) (Å) | 45.7 – 1.05 (1.08 – 1.05) |
| I/σI (high res. bin) | 18.92 (1.81) |
| R<sub>meas</sub> | 0.034 |
| CC(1/2) | 100 (66.6) |
| Resolution at CC(1/2) = 0.50 (Å) | 1.05 |
| Completeness (%) | 99.3 |
| Total observations | 817,928 |
| Unique observations | 284,505 |
| Redundancy | 2.9 |
| Wilson B-factor | 9.0 |
| Collected at beamline | ALS 12.3.1 |
| **Model Refinement** |  |
| Resolution (Å) | 45.7 – 1.05 |
| R<sub>work</sub>/R<sub>free</sub> | 0.130 / 0.146 |
| R<sub>work</sub>/R<sub>free</sub> at CC(1/2) = 0.5 res. cutoff | 0.130 / 0.146 |
| Mol. Per ASU (protein + ligand) | 3 |
| Number of atoms | 2975 |
| protein | 2378 |
| ligand/ions | 50 |
| waters | 547 |
| Average B-factor | 14.0 |
| r.m.s. deviations |  |
| Bond lengths (Å) | 0.014 |
| Bond angles (Å) | 1.588 |
| Ramachandran favored (%) | 97.83 |
| Ramachandran outliers (%) | 0.62 |
| All-atom clashscore | 2.23 |

## References

1. Florens L, Washburn MP, Raine JD, Anthony RM, Grainger M, Haynes JD, et al. A proteomic view of the Plasmodium falciparum life cycle. Nature. 2002;419(6906):520–6. Epub 2002/10/09. doi: 10.1038/nature01107. PubMed PMID: 12368866.

2. Royer RE, Deck LM, Campos NM, Hunsaker LA, Vanderjagt DL. Biologically-Active Derivatives of Gossypol - Synthesis and Antimalarial Activities of Peri-Acylated Gossylic Nitriles. Journal of medicinal chemistry. 1986;29(9):1799–801. doi: Doi 10.1021/Jm00159a043. PubMed PMID: ISI:A1986D792300043.

3. Boucher JI, Jacobowitz JR, Beckett BC, Classen S, Theobald DL. An atomic-resolution view of neofunctionalization in the evolution of apicomplexan lactate dehydrogenases. Elife. 2014;3. doi: 10.7554/eLife.02304. PubMed PMID: 24966208; PubMed Central PMCID: PMCPMC4109310.

4. Shoemark DK, Cliff MJ, Sessions RB, Clarke AR. Enzymatic properties of the lactate dehydrogenase enzyme from Plasmodium falciparum. Febs Journal. 2007;274(11):2738–48. doi: Doi 10.1111/J.1742-4658.2007.05808.X. PubMed PMID: ISI:000246681900006.

5. Gomez MS, Piper RC, Hunsaker LA, Royer RE, Deck LM, Makler MT, et al. Substrate and cofactor specificity and selective inhibition of lactate dehydrogenase from the malarial parasite P-falciparum. Molecular and Biochemical Parasitology. 1997;90(1):235–46. doi: Doi 10.1016/S0166-6851(97)00140-0. PubMed PMID: ISI:000071716100020.

6. Read JA, Wilkinson KW, Tranter R, Sessions RB, Brady RL. Chloroquine binds in the cofactor binding site of Plasmodium falciparum lactate dehydrogenase. Journal of Biological Chemistry. 1999;274(15):10213–8. doi: Doi 10.1074/Jbc.274.15.10213. PubMed PMID: ISI:000079663500045.

7. Brady RL, Cameron A. Structure-based approaches to the development of novel anti-malarials. Current drug targets. 2004;5(2):137–49. Epub 2004/03/12. PubMed PMID: 15011947.

8. Cameron A, Read J, Tranter R, Winter VJ, Sessions RB, Brady RL, et al. Identification and activity of a series of azole-based compounds with lactate dehydrogenase-directed anti-malarial activity. Journal of Biological Chemistry. 2004;279(30):31429–39. doi: Doi 10.1074/Jbc.M402433200. PubMed PMID: ISI:000222726800064.

9. Conners R, Schambach F, Read J, Cameron A, Sessions RB, Vivas L, et al. Mapping the binding site for gossypol-like inhibitors of Plasmodium falciparum lactate dehydrogenase. Molecular and Biochemical Parasitology. 2005;142(2):137–48. doi: Doi 10.1016/J.Molbiopara.2005.03.015. PubMed PMID: ISI:000230687000001.

10. Vivas L, Easton A, Kendrick H, Cameron A, Lavandera JL, Barros D, et al. Plasmodium falciparum: stage specific effects of a selective inhibitor of lactate dehydrogenase. Experimental parasitology. 2005;111(2):105–14. Epub 2005/08/16. doi: 10.1016/j.exppara.2005.06.007. PubMed PMID: 16098967.

11. Choi S-r, Pradhan A, Hammond NL, Chittiboyina AG, Tekwani BL, Avery MA. Design, Synthesis, and Biological Evaluation of Plasmodium falciparum Lactate Dehydrogenase Inhibitors. Journal of medicinal chemistry. 2007;50(16):3841–50. doi: 10.1021/jm070336k.

12. Granchi C, Bertini S, Macchia M, Minutolo F. Inhibitors of lactate dehydrogenase isoforms and their therapeutic potentials. Current medicinal chemistry. 2010;17(7):672–97. Epub 2010/01/22. PubMed PMID: 20088761.

13. Saxena S, Durgam L, Guruprasad L. Multiple e-pharmacophore modelling pooled with high-throughput virtual screening, docking and molecular dynamics simulations to discover potential inhibitors of Plasmodium falciparum lactate dehydrogenase (PfLDH). Journal of biomolecular structure & dynamics. 2018:1–17. Epub 2018/05/03. doi:10.1080/07391102.2018.1471417. PubMed PMID: 29718775.

14. Kaushal NA, Kaushal DC. Production and characterization of monoclonal antibodies against substrate specific loop region of Plasmodium falciparum lactate dehydrogenase. Immunological investigations. 2014;43(6):556–71. Epub 2014/04/08. doi:10.3109/08820139.2014.892962. PubMed PMID: 24702659.

15. Bork S, Yokoyama N, Ikehara Y, Kumar S, Sugimoto C, Igarashi I. Growth-Inhibitory Effect of Heparin on Babesia Parasites. Antimicrobial Agents and Chemotherapy. 2004;48(1):236–41. doi: 10.1128/AAC.48.1.236-241.2004. PubMed PMID: PMC310193.

16. Eventoff W, Rossmann MG, Taylor SS, Torff HJ, Meyer H, Keil W, et al. Structural Adaptations of Lactate-Dehydrogenase Isozymes. Proceedings of the National Academy of Sciences of the United States of America. 1977;74(7):2677–81. doi: Doi 10.1073/Pnas.74.7.2677. PubMed PMID: ISI:A1977DP86600023.

17. Guindon S, Dufayard JF, Lefort V, Anisimova M, Hordijk W, Gascuel O. New algorithms and methods to estimate maximum-likelihood phylogenies: assessing the performance of PhyML 3.0. Syst Biol. 2010;59(3):307–21. Epub 2010/06/09. doi:syq010 [pii] 10.1093/sysbio/syq010. PubMed PMID: 20525638.

18. Celniker G, Nimrod G, Ashkenazy H, Glaser F, Martz E, Mayrose I, et al. ConSurf: Using Evolutionary Data to Raise Testable Hypotheses about Protein Function. Israel Journal of Chemistry. 2013;53(3-4):199–206. doi: 10.1002/ijch.201200096.

19. Nicholls DJ, Miller J, Scawen MD, Clarke AR, Holbrook JJ, Atkinson T, et al. The importance of arginine 102 for the substrate specificity of Escherichia coli malate dehydrogenase. Biochem Biophys Res Commun. 1992;189(2):1057–62. Epub 1992/12/15. PubMed PMID: 1472016.

20. Ohshima T, Sakuraba H. Purification and characterization of malate dehydrogenase from the phototrophic bacterium, Rhodopseudomonas capsulata. Biochimica et Biophysica Acta (BBA) - Protein Structure and Molecular Enzymology. 1986;869(2):171–7. doi: https://doi.org/10.1016/0167-4838(86)90291-8.

21. Hartl T, Grossebuter W, Gorisch H, Stezowski JJ. Crystalline NAD/NADP-dependent malate dehydrogenase; the enzyme from the thermoacidophilic archaebacterium Sulfolobus acidocaldarius. Biological chemistry Hoppe-Seyler. 1987;368(3):259–67. Epub 1987/03/01. PubMed PMID: 3109450.

22. Wynne SA, Nicholls DJ, Scawen MD, Sundaram TK. Tetrameric malate dehydrogenase from a thermophilic Bacillus: cloning, sequence and overexpression of the gene encoding the enzyme and isolation and characterization of the recombinant enzyme. Biochemical Journal. 1996;317(Pt 1):235–45. PubMed PMID: PMC1217469.

23. Langelandsvik AS, Steen IH, Birkeland NK, Lien T. Properties and primary structure of a thermostable L-malate dehydrogenase from Archaeoglobus fulgidus. Archives of microbiology. 1997;168(1):59–67. Epub 1997/07/01. PubMed PMID: 9211715.

24. Madern D, Ebel C, Mevarech M, Richard SB, Pfister C, Zaccai G. Insights into the molecular relationships between malate and lactate dehydrogenases: structural and biochemical properties of monomeric and dimeric intermediates of a mutant of tetrameric L-[LDH-like] malate dehydrogenase from the halophilic archaeon Haloarcula marismortui. Biochemistry. 2000;39(5):1001–10. Epub 2000/02/02. PubMed PMID: 10653644.

25. Dando C, Schroeder ER, Hunsaker LA, Deck LM, Royer RE, Zhou X, et al. The kinetic properties and sensitivities to inhibitors of lactate dehydrogenases (LDH1 and LDH2) from Toxoplasma gondii: comparisons with pLDH from Plasmodium falciparum. Molecular and Bochemical Parasitology. 2001;118(23–32). doi:10.1016/S0166-6851(01)00360-7.

26. Madern D, Cai X, Abrahamsen MS, Zhu G. Evolution of Cryptosporidium parvum Lactate Dehydrogenase from Malate Dehydrogenase by a Very Recent Event of Gene Duplication. Molecular Biology and Evolution. 2004;21(3):489–97. doi: 10.1093/molbev/msh042.

27. Madern D. Molecular Evolution Within the L-Malate and L-Lactate Dehydrogenase Super-Family. Journal of Molecular Evolution. 2002;54(6):825–40. doi: 10.1007/s00239-001-0088-8.

28. Madern D, Cai XM, Abrahamsen MS, Zhu G. Evolution of Cryptosporidium parvum lactate dehydrogenase from malate dehydrogenase by a very recent event of gene duplication. Molecular Biology and Evolution. 2004;21(3):489–97. doi: Doi 10.1093/Molbev/Msh042. PubMed PMID: ISI:000220273600007.

29. Jeffery CJ. Moonlighting proteins. Trends in Biochemical Sciences. 1999;24(1):8–11. doi: https://doi.org/10.1016/S0968-0004(98)01335-8.

30. Huberts DHEW, van der Klei IJ. Moonlighting proteins: An intriguing mode of multitasking. Biochimica et Biophysica Acta (BBA) - Molecular Cell Research. 2010;1803(4):520–5. doi: https://doi.org/10.1016/j.bbamcr.2010.01.022.

31. Hendriks W, Mulders JW, Bibby MA, Slingsby C, Bloemendal H, de Jong WW. Duck lens epsilon-crystallin and lactate dehydrogenase B4 are identical: a single-copy gene product with two distinct functions. Proceedings of the National Academy of Sciences. 1988;85(19):7114.

32. Piatigorsky J. Multifunctional Lens Crystallins and Corneal Enzymes: More than Meets the Eye. Annals of the New York Academy of Sciences. 2006;842(1):7–15. doi: 10.1111/j.1749-6632.1998.tb09626.x.

